# Identification of a novel tedizolid resistance mutation in *rpoB* of methicillin-resistant Staphylococcus aureus

**DOI:** 10.1101/827501

**Authors:** Tianwei Shen, Kelsi Penewit, Adam Waalkes, Libin Xu, Stephen J. Salipante, Abhinav Nath, Brian J. Werth

## Abstract

A tedizolid-resistant isolate of MRSA was selected by serial passage. Whole genome sequencing revealed only a single nucleotide variant in *rpoB*. Cross-resistance to linezolid, chloramphenicol, and quinupristin-dalfopristin was observed but susceptibility to other drugs including rifampin was unchanged. Models of the RNA-polymerase-ribosomal complex revealed that the mutated residue was unlikely to interact directly with the oxazolidinone binding site. This is the first time that *rpoB* mutation has been associated with resistance to the PhLOPSa antimicrobials.

---

Tedizolid is an oxazolidinone antimicrobial with broad spectrum activity against Gram-positive bacteria including methicillin-resistant *S. aureus* (MRSA).^1^ Like other oxazolidinones tedizolid exerts its antibacterial activity by binding the 23s rRNA component of the 50s-ribosomal subunit and thus inhibiting protein synthesis. Resistance to tedizolid is uncommon but mutations in the ribosomal proteins L3, L4, and L22 (encoded by *rplC*, *rplD*, and *rplV* respectively), and the 23S rRNA target, which also mediate the so-called PhLOPSa (phenicol, lincosamide, oxazolidinone, pleuromutilin, and streptogramin A) resistance phenotype have been implicated.^2, 3^ Acquisition of the transferable rRNA methyltransferase gene, *cfr*, may also cause resistance to linezolid and other PhLOPSa antimicrobials but is generally believed to be insufficient to produce tedizolid resistance on its own even though it may increase the tedizolid MIC. The plasmid-carried *optrA* gene, which encodes an ABC transporter, has been implicated in oxazolidinone and phenicol resistance in enterococci and streptococci but has not been identified in *S. aureus*.^4^ Previous serial passage studies with tedizolid have seen limited success in selecting for tedizolid resistance and only mutations in the 23S rRNA have been recovered following tedizolid exposure.^5^ Genes affecting quinolone efflux such as *norA* and *mepA*, or lincosamide efflux such as *mdeA* have not been reported to affect oxazolidinone activity.

In this study, we observed the emergence of a mutant exhibiting a novel mechanism of PhLOPSa resistance from a well characterized MRSA strain after serial passage in escalating concentrations of tedizolid.

Using the well characterized MRSA strain, N315, we selected for tedizolid resistance by serial passage in escalating concentrations of tedizolid in Mueller Hinton II broth (MHB) starting with 0.5× the MIC. Once visible growth was observed a sample of the broth was diluted 1:1000 into fresh MHB with twice the previous concentration of tedizolid until an isolate with an MIC of ≥4 mg/mL was recovered. This MIC was selected since it is 1 log_2_ dilution above the breakpoint for resistance (MIC ≥ 2mg/L). After 10 passages, we recovered an isolate (N315-TDZ4) with a stable tedizolid MIC of 4mg/L or 16× the MIC of the parent strain, N315. To explore the cross resistance associated with this evolved strain we reevaluated susceptibility to a panel of other antimicrobial agents (Table 1) by broth microdilution in accordance with Clinical Laboratory Standards Institute (CLSI) guidelines^6^ or by gradient strip in the case of quinupristin-dalfopristin (Liofilchem®). Cross-resistance to chloramphenicol, linezolid, and quinupristin-dalfopristin was observed but susceptibility to other drugs tested was relatively unchanged (Table 1). N315 is resistant to clindamycin so lincosamide cross-resistance was unevaluable in this study and we did not have access to pleuromutilins, which are not yet a clinically important class of antimicrobials. While macrolide susceptibility is not considered part of the PhLOPSa group macrolides also target the 50s ribosome, however, erythromycin susceptibility was also unevaluable since N315 is resistant to erythromycin at baseline.

**Table 1.**
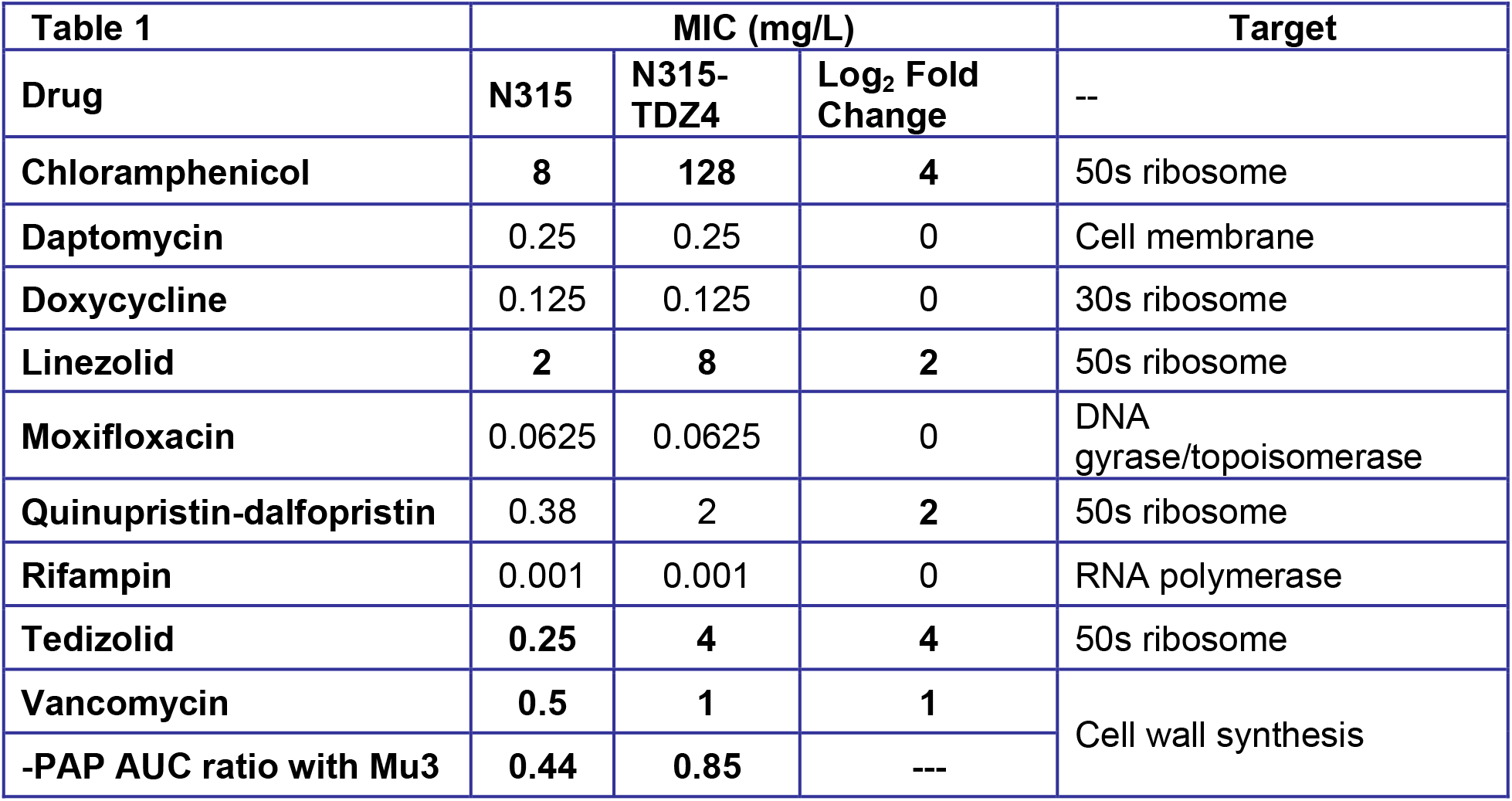
Minimum inhibitory concentrations (MIC) of parent strain, N315, and tedizolid passaged *rpoB* mutant, N315-TDZ4 to various antimicrobials, including fold change in MIC and antimicrobial target. PAP, population analysis profile; AUC, area under the cure

| Table 1 | MIC (mg/L) |  |  | Target |
| --- | --- | --- | --- | --- |
| Drug | N315 | N315-TDZ4 | Log<sub>2</sub> Fold Change | -- |
| Chloramphenicol | 8 | 128 | 4 | 50s ribosome |
| Daptomycin | 0.25 | 0.25 | 0 | Cell membrane |
| Doxycycline | 0.125 | 0.125 | 0 | 30s ribosome |
| Linezolid | 2 | 8 | 2 | 50s ribosome |
| Moxifloxacin | 0.0625 | 0.0625 | 0 | DNA gyrase/topoisomerase |
| Quinupristin-dalfopristin | 0.38 | 2 | 2 | 50s ribosome |
| Rifampin | 0.001 | 0.001 | 0 | RNA polymerase |
| Tedizolid | 0.25 | 4 | 4 | 50s ribosome |
| Vancomycin | 0.5 | 1 | 1 | Cell wall synthesis |
| -PAP AUC ratio with Mu3 | 0.44 | 0.85 | --- |  |

N315-TDZ4 was subjected to whole genome sequencing (WGS) using the MiSeq platform (Illumina, San Diego, CA, USA) as previously described^7^ to an average read depth of at least 50× per isolate and the. Sequence data from this study is freely available through the NCBI Sequence Read Archive (<u>PRJNA578164</u>). WGS of the N315-TDZ4 and comparison to the parent strain revealed a single nucleotide variant (SNV) in the *rpoB* gene (1345 A>G) corresponding to the amino acid substitution Asn449Asp. This mutation lies outside of the rifampin resistance-determining regions, which span from nucleotides 1384 – 1464 (AA 462–488) and 1543 – 1590 (AA 462–488) ^8, 9^, and we correspondingly did not observe any change in rifampin susceptibly (Table 1). Previous studies have reported an association between vancomycin and daptomycin susceptibility and certain rpoB mutations including Ala621Glu, and Ala477Asp, and His481Tyr.^10–12^ We did not observe any change in daptomycin MIC for our mutant but we did see a 1 log_2_-dilution increase in vancomycin MIC. To further assess the potential significance of this increase in vancomycin MIC we tested the N315-TDZ4 isolate and N315 parent strains for the heterogeneous vancomycin intermediate *S. aureus* (hVISA) phenotype by the gold standard population analysis profile (PAP) as previously described.^13^ Interestingly, we observed that PAP-AUC ratio with Mu3 of N315-TDZ4 was 0.85, up from 0.44 of the parent strain N315 (Table 1). While this increase is substantial it falls below the categorical criterion to declare this isolate an hVISA (PAP AUC ratio with Mu3 of ≥0.9).

In an effort to evaluate the potential impact of the rpoB Asn449Asp mutation on target protein function, we constructed a homology model of *S. aureus* rpoB using I-TASSER,^14^ and modeled the relative orientations of the RNA polymerase (RNAP) and the ribosome (Figure 1). This model suggests that the RNAP mutation would be unlikely to interact directly with binding sites for the oxazolidinones and other PhLOPSa drugs located in the 50S ribosomal subunit (Figure 1). This finding suggests an indirect mechanism of resistance, possibly involving one of the many factors that interact with the beta-subunit of RNAP.^8^

**Figure 1.**
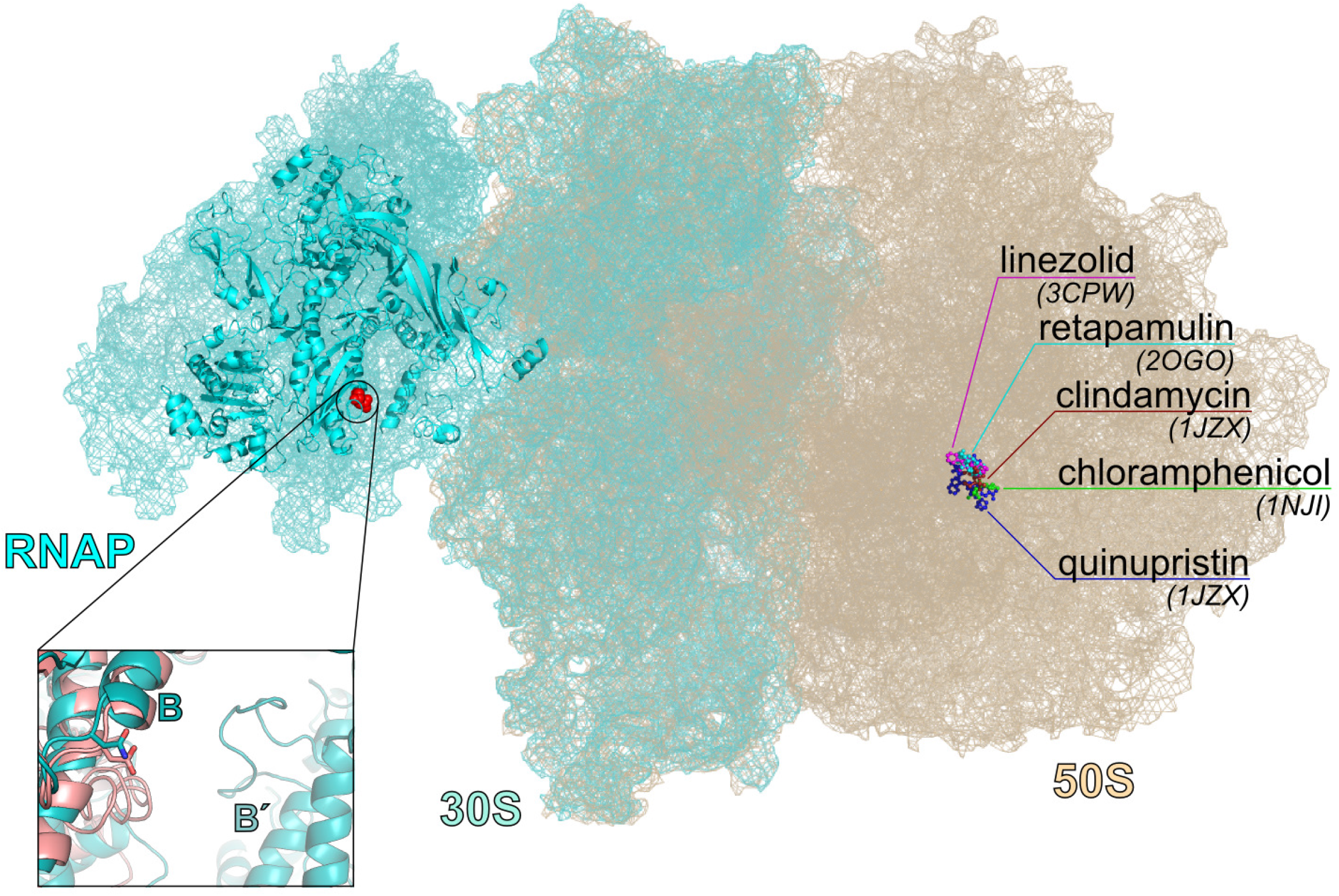
Model of *S. aureus* rpoB N449D docked into the RNA polymerase (RNAP)-ribosome transcriptional complex (protein data bank, IDs 5MY1 and 5U9F), illustrating that the mutated residue in RNAP lies ~170Å from the binding site of PHLOPSa drugs on the 50S ribosomal subunit. Drug binding sites are drawn from the PDB IDs indicated in parentheses. The great distance between the mutated residue and linezolid binding site suggests an indirect mechanism; notably, the susceptibility of doxycycline, which acts on the 30S ribosomal subunit, was not affected. One CryoEM study of the structure of the RNAP-ribosomal complex indicates an S1 protein crosslink between the 30S and RNAP at the helix-turn-helix affected by this mutation.

In this study, we report for the first time a novel mechanism of resistance to oxazolidinones, phenicols, and streptogramins involving mutation in *rpoB*, which encodes the β-subunit of RNAP. The precise molecular mechanism by which this mutation mediates this resistance phenotype is unclear but may involve transcriptional modulation by altered sigma-factor binding. Previous studies have demonstrated that the 30s-ribosomal protein S10 encoded by rpsJ links directly with the β-subunit of RNAP facilitating the tight linkage between transcription and translation in bacteria but the RNAP only interacts with the leading 30s-subunit and no known interaction between the 50s subunits and the RNAP exist. The fact that cross resistance was isolated to 50s-ribosomally active agents and not 30s active agents (doxycycline) suggests a relatively specific interaction with the 50s-ribosome and not a broader modulation of translation or protein synthesis. Based on the absence of altered susceptibility to other intracellularly active agents including moxifloxacin and doxycycline it seems unlikely that this variant facilitates multidrug efflux. For now, tedizolid resistance remains uncommon clinically and it is unknown whether this unique mutation could be selected for by other PhLOPSa drugs or would be likely to emerge after clinical exposure to tedizolid.

## Acknowledgments

This study was initially presented in part at the annual ID-Week conference in San Francisco, CA 2018. This study was supported in part by National Institute of Allergy and Infectious Diseases of the National Institutes of Health under the award numbers 1R21AI132994-01A1, 1R01AI136979-01 and by the University of Washington.

## References

1. Kisgen JJ, Mansour H, Unger NR, Childs LM. Tedizolid: a new oxazolidinone antimicrobial. American journal of health-system pharmacy : AJHP : official journal of the American Society of Health-System Pharmacists. 2014;71(8):621–33. doi: 10.2146/ajhp130482. PubMed PMID: 24688035.

2. Shaw KJ, Poppe S, Schaadt R, Brown-Driver V, Finn J, Pillar CM, Shinabarger D, Zurenko G. In vitro activity of TR-700, the antibacterial moiety of the prodrug TR-701, against linezolid-resistant strains. Antimicrob Agents Chemother. 2008;52(12):4442–7. doi: 10.1128/AAC.00859-08. PubMed PMID: 18838596; PMCID: PMC2592863.

3. Freitas AR, Dilek AR, Peixe L, Novais C. Dissemination of Staphylococcus epidermidis ST22 With Stable, High-Level Resistance to Linezolid and Tedizolid in the Greek-Turkish Region (2008-2016). Infect Control Hosp Epidemiol. 2018;39(4):492–4. doi: 10.1017/ice.2018.5. PubMed PMID: 29427999.

4. Wang Y, Lv Y, Cai J, Schwarz S, Cui L, Hu Z, Zhang R, Li J, Zhao Q, He T, Wang D, Wang Z, Shen Y, Li Y, Fessler AT, Wu C, Yu H, Deng X, Xia X, Shen J. A novel gene, optrA, that confers transferable resistance to oxazolidinones and phenicols and its presence in Enterococcus faecalis and Enterococcus faecium of human and animal origin. J Antimicrob Chemother. 2015;70(8):2182–90. doi: 10.1093/jac/dkv116. PubMed PMID: 25977397.

5. Locke JB, Hilgers M, Shaw KJ. Novel ribosomal mutations in Staphylococcus aureus strains identified through selection with the oxazolidinones linezolid and torezolid (TR-700). Antimicrob Agents Chemother. 2009;53(12):5265–74. doi: 10.1128/AAC.00871-09. PubMed PMID: 19752277; PMCID: PMC2786364.

6. Clinical and Laboratory Standards Institute. Performance Standards for Antimicrobial Susceptibility Testing: Twenty-fifth Informational Supplement M100-S27 CLSI, Wayne, PA. USA2017.

7. Roach DJ, Burton JN, Lee C, Stackhouse B, Butler-Wu SM, Cookson BT, Shendure J, Salipante SJ. A Year of Infection in the Intensive Care Unit: Prospective Whole Genome Sequencing of Bacterial Clinical Isolates Reveals Cryptic Transmissions and Novel Microbiota. PLoS Genet. 2015;11(7):e1005413. doi: 10.1371/journal.pgen.1005413. PubMed PMID: 26230489; PMCID: PMC4521703.

8. Alifano P, Palumbo C, Pasanisi D, Tala A. Rifampicin-resistance, rpoB polymorphism and RNA polymerase genetic engineering. J Biotechnol. 2015;202:60–77. doi: 10.1016/j.jbiotec.2014.11.024. PubMed PMID: 25481100.

9. Zhou W, Shan W, Ma X, Chang W, Zhou X, Lu H, Dai Y. Molecular characterization of rifampicin-resistant Staphylococcus aureus isolates in a Chinese teaching hospital from Anhui, China. BMC microbiology. 2012;12:240. doi: 10.1186/1471-2180-12-240. PubMed PMID: 23082766; PMCID: PMC3485161.

10. Cui L, Isii T, Fukuda M, Ochiai T, Neoh HM, Camargo IL, Watanabe Y, Shoji M, Hishinuma T, Hiramatsu K. An RpoB mutation confers dual heteroresistance to daptomycin and vancomycin in Staphylococcus aureus. Antimicrob Agents Chemother. 2010;54(12):5222–33. doi: 10.1128/AAC.00437-10. PubMed PMID: 20837752; PMCID: PMC2981288.

11. Baek KT, Thogersen L, Mogenssen RG, Mellergaard M, Thomsen LE, Petersen A, Skov S, Cameron DR, Peleg AY, Frees D. Stepwise decrease in daptomycin susceptibility in clinical Staphylococcus aureus isolates associated with an initial mutation in rpoB and a compensatory inactivation of the clpX gene. Antimicrob Agents Chemother. 2015;59(11):6983–91. doi: 10.1128/AAC.01303-15. PubMed PMID: 26324273; PMCID: PMC4604412.

12. Watanabe Y, Cui L, Katayama Y, Kozue K, Hiramatsu K. Impact of rpoB mutations on reduced vancomycin susceptibility in Staphylococcus aureus. J Clin Microbiol. 2011;49(7):2680–4. doi: 10.1128/JCM.02144-10. PubMed PMID: 21525224; PMCID: PMC3147882.

13. Wootton M, Howe RA, Hillman R, Walsh TR, Bennett PM, MacGowan AP. A modified population analysis profile (PAP) method to detect hetero-resistance to vancomycin in Staphylococcus aureus in a UK hospital. J Antimicrob Chemother. 2001;47(4):399–403. Epub 2001/03/27. PubMed PMID: 11266410.

14. Yang J, Zhang Y. Protein Structure and Function Prediction Using I-TASSER. Curr Protoc Bioinformatics. 2015;52:5 8 1–15. doi: 10.1002/0471250953.bi0508s52. PubMed PMID: 26678386; PMCID: PMC4871818.

